# Pathogen context reshapes antimicrobial peptide generation

**DOI:** 10.64898/2026.07.01.735178

**Authors:** Shiyang You, Chen Zhang, Yingchun Han, Qiuyun Jiang, Xuan Guo, Mingjia Li, Yongxin Su, Xiyang Dong, Menglin Yang, Hongyuan Lu

## Abstract

Antimicrobial peptides are a promising source of new anti-infective agents, but their discovery is limited by a practical bottleneck: selecting a small set of candidates to synthesize and test for a specific pathogen. Existing peptide generators can expand antimicrobial-like sequence space, but most leave pathogen specificity to predictors or filters after generation. Here, we use AMPHORA, a pathogen-conditioned sequence–structure generator, to build antimicrobial peptide libraries shaped by target context. AMPHORA couples a short-peptide representation model to latent-flow generation conditioned on target class, genome-derived features and strain-description text. In matched, partial and shuffled controls, pathogen context redirected generated peptide pools beyond broad activity labels, with the strongest effects when genome and text conditions were combined. Counterfactual generation from the same initial noise showed that strain descriptions primarily changed amino-acid choices, whereas genome-derived features contributed more strongly to predicted structural properties. Across species, AMPHORA produced target-dependent enrichment, and matched bacterial context shifted APEX-predicted potency-score distributions compared with class-only generation. The generated peptides remained diverse, novel and peptide-like by sequence and predicted-structure analyses. Collectively, our results establish pathogen context as an active design signal for antimicrobial peptide generation and provide a target-aware route to pre-synthesis library design.

## Introduction

Antimicrobial peptides are a promising source of new anti-infective molecules^[1]^. Machine-learning screens have expanded the accessible AMP search space across large biological sequence collections^[2]^. A major bottleneck arises before activity is measured: a laboratory must choose a small set of candidates to synthesize and test for a given pathogen. This choice is limited by synthesis capacity, assay throughput and the cost of follow-up characterization. Because AMP activity and liability profiles are target and assay dependent, a library suited to *Acinetobacter baumannii* may be poorly matched to *Candida albicans*, *Pseudomonas aeruginosa* or *Staphylococcus epidermidis*^[3]^. A generator that treats these targets as interchangeable can expand sequence space without improving the pool chosen for the assay.

AMPs make this selection problem especially difficult. These molecules are often short, charged and amphipathic; many disrupt microbial membranes^[4]^, and others act through intracellular or immunomodulatory mechanisms^[1]^. Small changes in residue composition or patterning can shift potency, solubility, selectivity, haemolysis, mammalian cytotoxicity and serum stability^[3]^. This sensitivity makes AMPs attractive design substrates and complicates pathogen-specific selection. A generator can produce plausible AMP-like sequences and still fail to bias the candidate pool toward the pathogen being tested.

Most generative AMP workflows generate first and address pathogen specificity later. Deep generative models can produce potent AMP-like sequences^[5]^, and recent generative AI pipelines have further broadened de novo AMP discovery^[6]^. They produce sequences with broad antimicrobial features, then use predictors, filters or manual rules to select candidates for synthesis^[3]^. By then, the candidate library has already been formed. For pathogen-specific discovery, the target organism should shape the candidate pool as it is made.

A pathogen target is more than an activity label. Broad labels such as antibacterial, antifungal, antiviral and antiparasitic distinguish major activity classes; they do not define a species or strain. Genome-derived features can encode organism-level sequence composition and strain-associated properties. Strain descriptions can capture taxonomy, source, phenotype and resistance-associated context. Together, these inputs bring target-organism information into peptide generation.

Here, we develop AMPHORA to test whether pathogen context can guide antimicrobial peptide generation before screening. AMPHORA couples a short-peptide representation backbone to a sequence–structure latent-flow generator that uses target class, genome-derived features and strain-description text^[7;8]^. The central question is whether these inputs change the candidate pool produced for a target while preserving the diversity, novelty and peptide-like properties needed for experimental follow-up.

We first established a decodable short-peptide generative backbone, then introduced pathogen inputs under matched, partial and shuffled conditions. Same-noise counterfactuals separated genome and text effects, species-level analyses measured target dependence, and novelty, diversity and predicted-structure analyses assessed whether these target-shaped libraries remained suitable for follow-up.

## Results

### A decodable short-peptide generator anchors the design space

Before introducing pathogen context, we first established a generator whose outputs could be evaluated as short-peptide candidates. Short AMPs and smORF-derived peptides concentrate functional information in local motifs, charge placement, hydrophobic and polar patterning, amphipathicity and partial disorder. A peptide-focused representation was needed before target information could be tested as a generative signal.

The peptide branch of AMPHORA combines a short-peptide encoder, deterministic biophysical priors, decodable latent compression and a sequence–structure latent-flow generator (Fig. 1). This separated baseline peptide generation from the pathogen-conditioning tests. Peptide hidden states were mapped into a 256-dimensional latent sequence and decoded back into canonical amino-acid strings, producing valid outputs. Conditional effects could therefore be evaluated directly in generated peptides.

**Figure 1.**
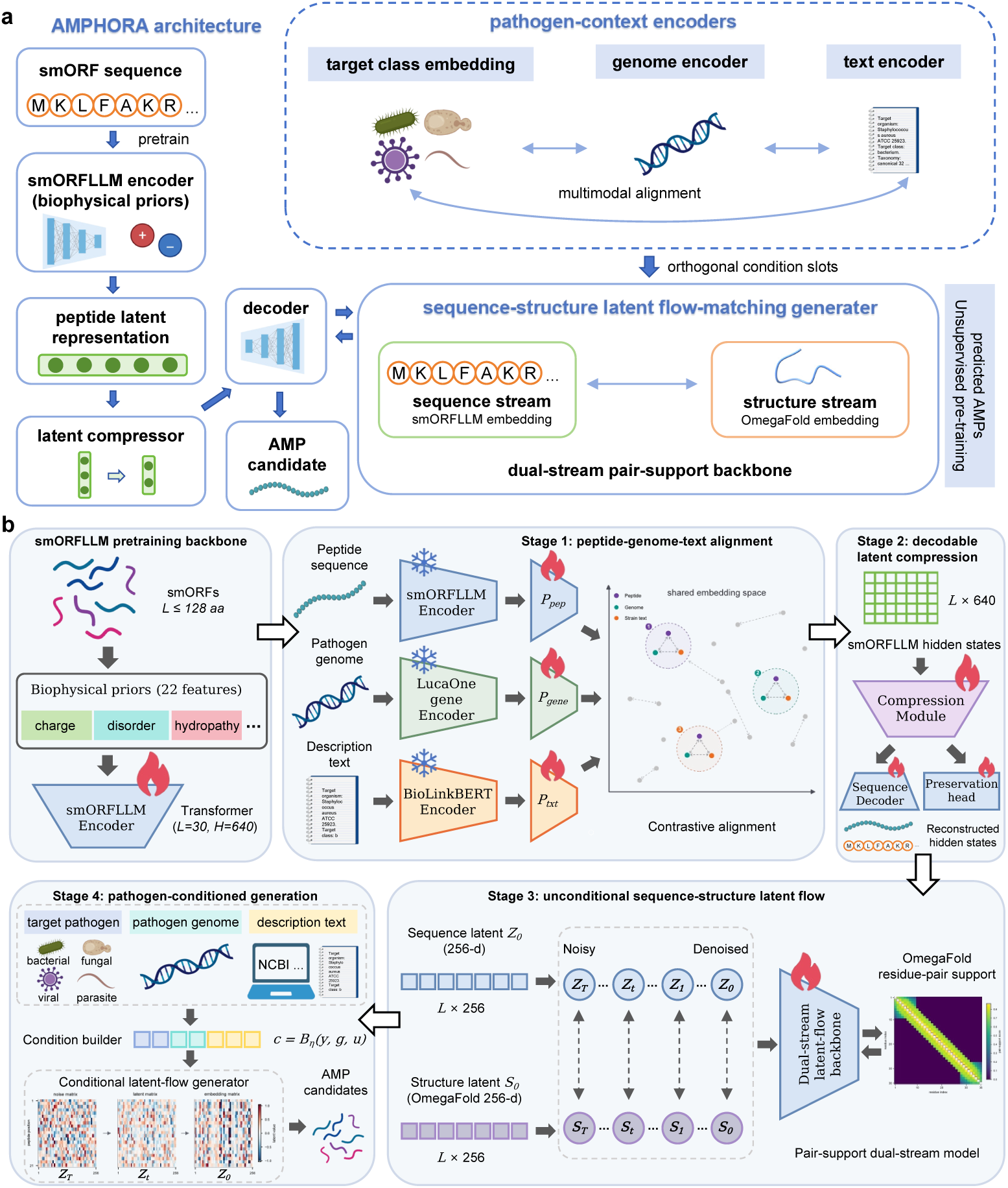
AMPHORA architecture for pathogen-context-conditioned antimicrobial peptide generation. **a,** Overview of the AMPHORA framework. A short-peptide or smORF sequence is encoded by smORFLLM, a peptide-focused encoder augmented with deterministic biophysical priors, to produce peptide latent representations that remain decodable through a learned latent compressor and sequence decoder. In parallel, pathogen context is represented through three aligned inputs: target-class embeddings, genome-derived features and strain-description text embeddings. These pathogen-context encoders are aligned with peptide representations and injected as orthogonal condition slots into a sequence–structure latent flow-matching generator. The generator uses a dual-stream pair-support backbone, with a sequence stream derived from smORFLLM embeddings and a structure stream derived from OmegaFold embeddings, to produce pathogen-context-shaped antimicrobial peptide libraries for downstream computational evaluation and experimental screening. **b,** Training and generation stages. The smORFLLM pretraining backbone learns short-peptide representations from smORF and small-protein sequences capped at 128 amino acids, with biophysical priors including charge, disorder and length-related descriptors. In Stage 1, peptide sequences, pathogen genomes and strain-description texts are encoded by smORFLLM, LucaOne and BioLinkBERT, respectively, and projected into a shared peptide-conditioned embedding space using contrastive alignment. In Stage 2, smORFLLM residue-level hidden states are compressed into a 256-dimensional decodable latent sequence; a sequence decoder preserves amino-acid reconstructability, whereas a preservation head retains information from the original smORFLLM hidden states. In Stage 3, an unconditional sequence–structure latent flow is trained over sequence and OmegaFold-derived structure latents, with residue-pair support used to select the dual-stream pair-support backbone. In Stage 4, target pathogen class, genome features and description text are assembled by a condition builder, *c* = *B_η_* (*y*, *g*, *u*), and used to guide conditional latent-flow generation from noise to peptide latent states, yielding pathogen-context-shaped AMP candidates.

We next compared four unconditional flow backbones under the same evaluation setting: sequence-only, global-structure latent, joint sequence–structure and pair-support dual-stream models (Fig. 2a). The pair-support dual-stream model emerged as the most balanced backbone, preserving local peptide grammar and predicted structural features. In the fixed-25-amino-acid benchmark, it achieved PRDC precision 0.934, PRDC recall 0.403, uniqueness 1.0000 and an APEX-predicted MIC < 100 *μ*M fraction of 0.070 (Fig. 2b and Supplementary Table 5). This benchmark selected a backbone that could generate short, valid and AMP-like candidates before any pathogen information was introduced. The pair-support dual-stream backbone was carried forward into the pathogen-conditioned generation experiments.

**Figure 2.**
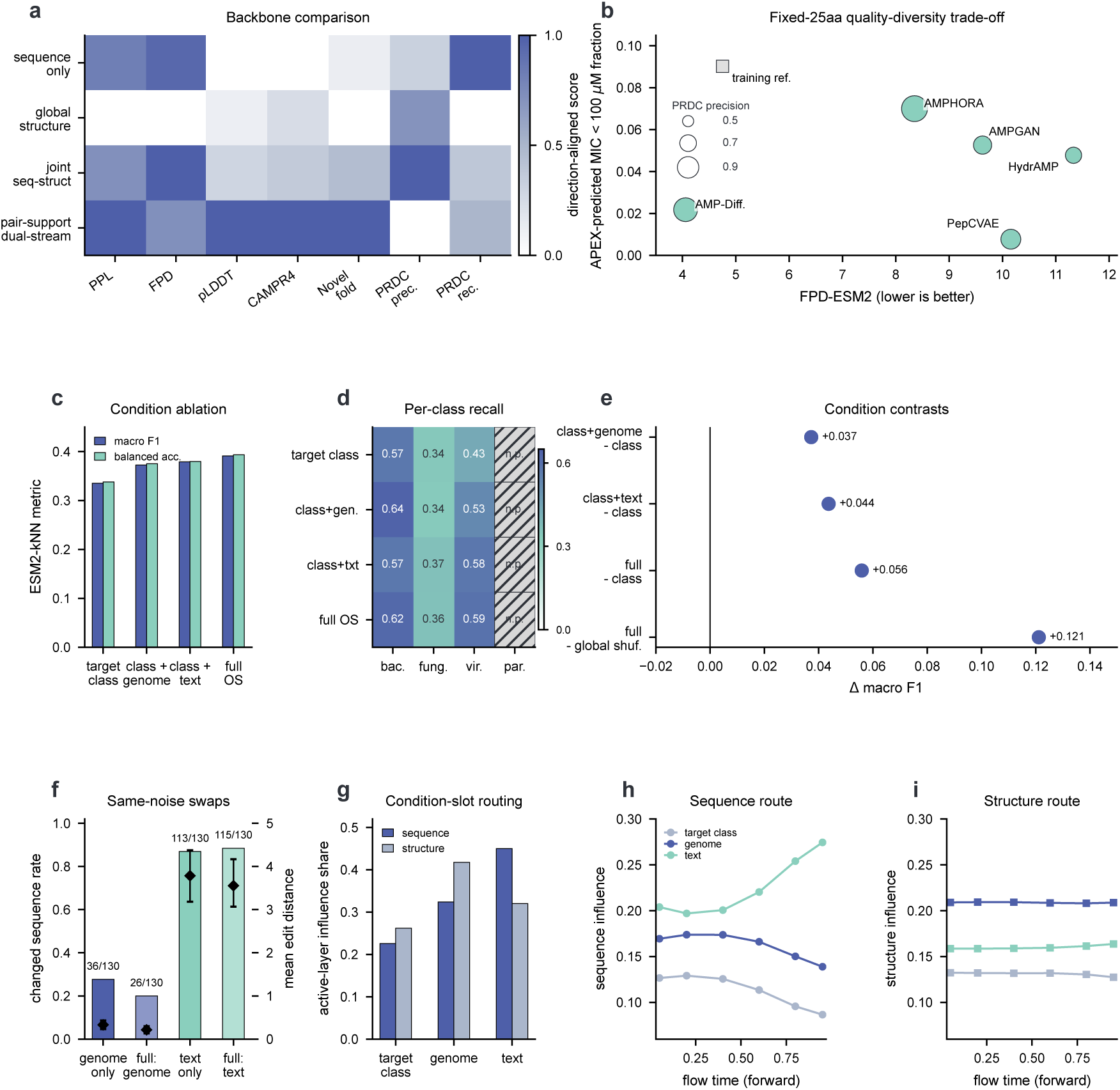
Backbone selection and pathogen-context controls. **a,** Retained Stage 3 backbones compared across direction-aligned metrics; darker cells indicate stronger scaled performance. Pseudo-perplexity measures peptide-sequence grammar, FPD measures embedding-distribution distance to reference peptides, pLDDT measures predicted structural confidence, CAMP_R4_ measures AMP-predictor support, and PRDC precision and recall measure local fidelity and coverage. **b,** Fixed-25-amino-acid quality–diversity trade-off under common output-level re-evaluation. The y-axis is the fraction of peptides with APEX-predicted MIC below 100 *μ*M. Marker size encodes PRDC precision. The training reference is shown as an anchor and was not ranked as a generator. **c,** Partial-condition comparison for target class only, target class plus genome, target class plus text and full orthogonal-slot conditioning. **d,** Per-class recall for the same partial-condition settings. Parasite cells are marked n.p. because validation support was limited. **e,** Observed macro F1 contrasts computed from the same final Stage 4 condition and shuffling controls used in Supplementary Tables 6 and 11. **f,** Same-noise counterfactual swaps with target class, length, latent noise, solver and seed fixed. Bars show changed-sequence rate, labels give changed rows out of 130, and diamonds show mean edit distance with 95% confidence intervals. **g,** Active-layer influence share for target-class, genome and text slots in sequence and structure streams. **h, i,** Sequence- and structure-stream influence over flow time. Flow time is shown in forward-time coordinates; sampling proceeds from *t* = 1 to *t* = 0.

### Pathogen context redirects generated peptide pools beyond target class

We next tested whether pathogen context redirected generated peptide pools beyond target class. Target-class-only generation provided the baseline, as antibacterial, antifungal, antiviral and antiparasitic labels already organize AMP sequence space. These labels do not specify the species or strain used in an assay. We compared matched full conditioning with target-class-only, class-plus-genome, class-plus-text and shuffled genome/text conditions. Full conditioning increased macro F1 from 0.335 with target class alone to 0.391 (Fig. 2c and Supplementary Table 6). Target class plus genome reached 0.373, and target class plus text reached 0.379. Genome and text each improved assignment beyond target class, and the full condition gave the strongest signal. This ordering placed pathogen-derived information above the broad class label and showed that both modalities contributed to the generated pool.

Shuffling genome and text conditions weakened this signal. Global reassignment reduced macro F1 to 0.270, below the target-class-only baseline (Fig. 2e and Supplementary Table 11). The drop indicates that the matched signal was not explained by a generic AMP prior or target-class labels alone. Within-class shuffling stayed closer to matched generation, showing that part of the readout follows broad class structure in the paired corpus. Thus, pathogen context added information beyond class labels, while broad activity class remained a major organizer of AMP space.

Class-wise recall showed detectable bacterial, fungal and viral signals; parasite support was too sparse for a class-level conclusion (Fig. 2d). Thus, aligned pathogen context altered the generated pool before any post-generation filtering or external scoring.

### Genome and text conditions shape different parts of generation

Genome and text conditions changed generation in different ways. We isolated each channel by generating paired peptides with the same target class, length, initial noise, solver settings and seed, then swapping either the genome or text condition. Paired outputs differed only in the exchanged condition channel. This same-noise design allowed sequence changes to be attributed to the swapped pathogen input.

Text-swap conditions changed 113–115 of 130 paired outputs and produced the larger edit distances (Fig. 2f). Genome-swap conditions changed 26–36 of 130 paired outputs and produced sparser sequence edits. Strain-description text was the stronger sequence-level driver under matched sampling conditions.

We then asked whether these output-level differences were reflected in the internal condition routes of the flow model. Condition-slot analyses separated the effects across the sequence and structure streams (Fig. 2g–i). Text contributed most strongly to the sequence stream; genome-derived features contributed more to the structure stream. Across flow time, text influence increased along the sequence stream, and genome influence remained higher along the structure stream. These trends were consistent with the same-noise swaps: text conditions produced larger amino-acid changes, and genome-derived features acted more strongly through the predicted-structure path.

Strain-description text mainly altered amino-acid choices, whereas genome-derived features acted more strongly on predicted structural properties.

### Pathogen responsiveness is target dependent

Antimicrobial screens ultimately ask whether candidates work against a species or strain. We next tested whether matched conditioning enriched generated pools at the species level. This readout is more demanding than class assignment because species reference sets are smaller and less balanced. It also tests the central claim more directly: a pathogen-conditioned generator should change candidate pools for individual targets, not only for broad AMP classes.

Each species was represented by multiple peptide prototypes, reducing dependence on a single reference centroid. Fresh matched candidates were generated for 12 species, with three seeds and 1,000 peptides per species and seed. Across the panel, mean diagonal enrichment was 0.00613 and the mean specificity ratio was 1.075. The pooled effect was small, but the target-by-target pattern was informative. *Candida albicans* (0.0187), *Aspergillus fumigatus* (0.0146) and *Staphylococcus epidermidis* (0.0112) showed the strongest matched enrichment. *Klebsiella pneumoniae* fell below baseline (-0.0067) (Fig. 3a).

**Figure 3.**
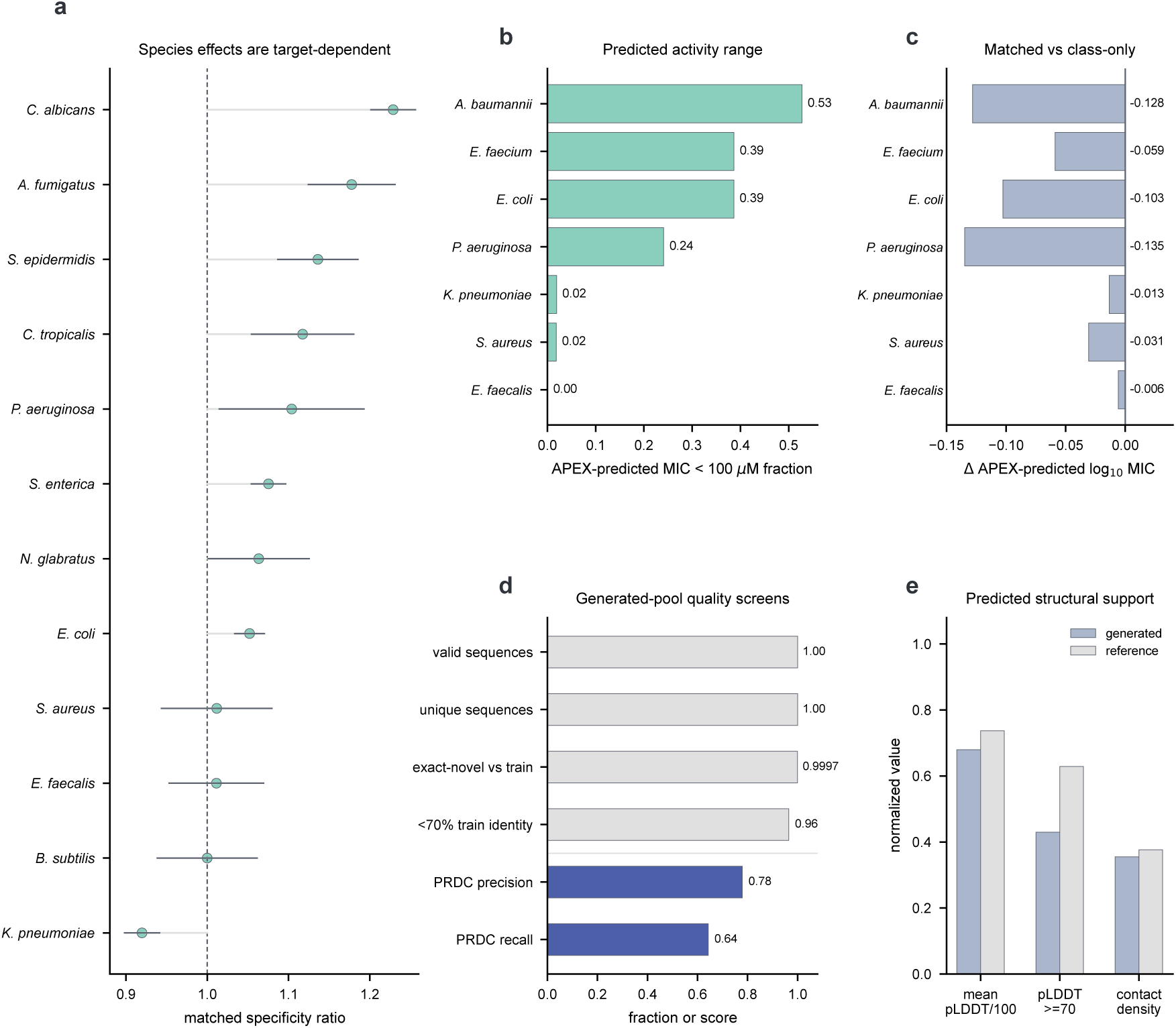
Organism-aware generation produces enriched and usable candidate pools. **a,** Per-species matched specificity ratios in the 12-organism enrichment benchmark. Points show means across three seeds, horizontal lines show 95% confidence intervals, and the dashed line marks ratio 1. **b,** Fraction of matched generated peptides with species-level APEX-predicted MIC below 100 *μ*M across seven bacterial species. **c,** Difference in species-level APEX-predicted log_10_ MIC between matched and target-class-only generation; negative values indicate lower predicted MIC under matched conditioning. **d,** Sequence novelty, uniqueness and AMP-neighbourhood support for matched AMPHORA-OS generation. **e,** Generated-versus-reference structural source metrics for the final Stage 4 structure panel. APEX scores are external predicted-activity estimates, not measured MIC values.

This pattern showed that species-level enrichment was target dependent across the 12-species panel.

We then asked whether this species-level shift also affected an external predicted-potency readout. Matched and target-class-only bacterial libraries were evaluated with APEX, a species-level antimicrobial potency predictor. Across seven bacterial species, the fraction of matched generated peptides with APEX-predicted MIC below 100 *μ*M ranged from 0.528 for *Acinetobacter baumannii* to 0.00 for *Enterococcus faecalis* (Fig. 3b). Matched generation produced lower species-level APEX-predicted log_10_ MIC than target-class-only generation for all seven species (Fig. 3c). The largest shifts were observed for *Pseudomonas aeruginosa* (-0.135), *Acinetobacter baumannii* (-0.128) and *Escherichia coli* (-0.103). Thus, matched bacterial context shifted the distribution of APEX-predicted potency scores, in addition to species-level peptide-space enrichment.

### Conditioned libraries retain novelty and peptide-like support

We next evaluated whether target-shaped libraries retained sequence novelty and peptide-like support. The analysis focused on validity, uniqueness, training-set similarity, AMP-neighbourhood support and predicted structural features.

Matched full-condition generation produced valid, unique peptide sequences. Exact-match novelty against the training set was 0.9997, and 96.4% of generated peptides remained below 70% nearest-training identity (Fig. 3d). These two measures separate exact memorization from near-neighbour similarity. Generated peptides also retained local AMP-reference support without concentrating in a narrow training-like cluster (Fig. 3d). The conditioned libraries therefore remained near AMP-like regions while retaining substantial novelty relative to the training set.

We next tested whether this sequence-level novelty was accompanied by predicted structural support. OmegaFold-derived models had a mean pLDDT of 67.95, compared with 73.72 for curated references, and contact density remained close to the reference panel (Fig. 3e and Supplementary Table 10). A lineage-level comparison showed higher predicted structural confidence, lower disorder and higher contact density in the final AMPHORA with orthogonal condition slots (AMPHORA-OS) checkpoint than in the no-condition Stage 3 predecessor (Supplementary Table 9). Pathogen conditioning preserved the sequence–structure regime selected during backbone development.

Pathogen-context shifts therefore occurred within a novel, peptide-like design space. They were not driven by copying, invalid decoding or motif collapse. The resulting libraries provided diverse, novel and peptide-like candidate pools for computational evaluation and experimental follow-up.

## Discussion

AMPHORA moves pathogen specificity from post-generation filtering into the generative step of antimicrobial peptide design. The central bottleneck is not simply whether a model can produce AMP-like sequences, but how the candidate-library distribution should be shaped for a defined pathogen under a finite experimental budget. By conditioning generation on target class, genome-derived features and strain-description text, AMPHORA makes pathogen context part of library construction. This reframes AMP generation as a pre-synthesis allocation problem in which the target organism shapes the peptide pool sampled before screening.

The evidence supports a controlled computational claim. A decodable short-peptide generator first established a peptide-like design space in which conditional effects could be measured. Matched, partial and shuffled controls then showed that aligned pathogen inputs changed generated peptide pools beyond broad activity labels. Same-noise counterfactuals and condition-slot analyses separated the two conditioning routes: strain-description text mainly altered amino-acid choices, whereas genome-derived features acted more strongly through predicted structural properties. Species-level analyses extended the readout beyond class labels and showed target-dependent enrichment across pathogens. Novelty, identity, AMP-neighbourhood and predicted-structure checks showed that these shifts occurred within a diverse, non-memorized and peptide-like candidate space.

This contribution differs from scaffold optimization and reservoir mining. Scaffold optimization begins with a known peptide and searches locally for improved variants. Reservoir mining searches existing biological sequence space for candidates that are already present. AMPHORA instead asks how a target organism should shape de novo candidate-library construction before synthesis. This distinction matters because antimicrobial discovery is often limited by assay capacity. A computational generator is useful only if it changes the composition of the small library that will actually be tested.

The uneven species-level response defines the operating range of the current model. Matched context produced clearer enrichment for some pathogens than for others, and within-class shuffling showed that broad activity labels still explain part of the generated signal. These results place AMPHORA at the level of candidate-pool allocation: it uses pathogen-associated genome and text information to shape which peptides enter a finite design set before synthesis. Predicted structural analyses further show that this reshaping occurs within a peptide-like sequence–structure regime, while leaving activity, selectivity and mechanism to experimental screening.

This positioning suggests a direct path for experimental use. Matched, mismatched and target-class-only libraries can be synthesized under the same budget and compared across pathogens with strong and weak computational enrichment. The relevant outcome is whether matched pathogen context allocates the same synthesis effort to a better candidate set, measured by hit rate, potency, selectivity and developability-associated properties. AMPHORA therefore positions pathogen context as a species-aware generative signal, shifting the pre-synthesis question from which individual peptide to rank first to what target-shaped candidate library should be sampled under a fixed experimental budget.

## Methods

### Study design and training stages

AMPHORA was trained through four stages with separate objectives. Stage 1 aligned peptide, genome and strain-description text representations on a curated conditional antimicrobial peptide (AMP) corpus. Stage 2 compressed peptide hidden states into a decodable 256-dimensional latent sequence. Stage 3 trained unconditional latent-flow generators and selected the sequence–structure backbone used for conditional modelling. Stage 4 introduced target-class, genome and text conditions through orthogonal condition slots and evaluated matched, partial and shuffled condition settings. Unless otherwise stated, peptide sequences used the 20 standard amino acids and were capped at 128 residues. Detailed objectives, optimization settings and checkpoint-selection rules are provided in Supplementary Methods.

### Peptide and pathogen-condition corpora

The peptide representation model was pretrained on a short-peptide/smORF corpus rather than on AMP activity labels. Preprocessing started from a representative FASTA containing 390,709,416 sequences assembled from public short-protein/smORF resources and internal smORF collections. MMseqs2 cluster-consistent splitting at 50% sequence identity produced 386,797,786 training rows, 1,951,071 validation rows and 1,960,384 test rows^[9]^. Length-aware sampling retained all sequences of length at most 50, sequences of length 51–90 with probability 0.5 and longer sequences with probability 0.2.

The conditional corpus was derived from the AMP-positive subset of ESCAPE, a standardized multilabel AMP benchmark integrating more than 80,000 peptides from 27 validated repositories^[10]^. The ESCAPE-derived AMP-positive source table contained 21,409 unique AMP sequences. Peptide–target pathogen conditions were constructed by combining dbAMP and DRAMP target annotations with ESCAPE category labels, followed by manual pathogen/category checks. Exact dbAMP or DRAMP matches were treated as target-annotation evidence. Records supported only by ESCAPE category fallback labels were retained as target-class or category-level conditions rather than validated species-specific activity evidence.

The formal conditional split contained 93,834 training rows and 4,910 validation rows, corresponding to 20,313 and 1,070 unique peptides. Exact train–validation peptide-sequence overlap was zero. The training set contained 73,500 bacterial, 13,745 fungal, 5,999 viral and 590 parasite rows. The validation set contained 3,842 bacterial, 833 fungal, 215 viral and 20 parasite rows. These counts define the long-tailed structure retained in the prespecified four-class analyses.

### Genome and text condition processing

Manually checked conditions were expanded into peptide–pathogen–genome–text records. Each retained row was assigned a canonical species or taxon identifier, a representative genome cache key and a standardized strain-description text template through a manually curated pathogen mapping table. Genome and text conditions were cached before conditional generation and paired to peptide–pathogen records through these cache keys.

Genome embeddings were produced with LucaOne gene encoder checkpoint LucaGroup/LucaOne-gene-step36.8M^[7]^. Representative genome FASTA files were split into windows of up to 4,096 nt with a 2,048-nt stride, and up to 64 windows were retained by uniform, high-complexity and deterministic random sampling. LucaOne window embeddings were mean-pooled to a 2,560-dimensional raw genome representation. The exported Stage 1 genome adapter projected this representation into the 256-dimensional AMPHORA condition space.

Text embeddings were produced from whitespace-normalized strain-description templates with BioLinkBERT-large checkpoint michiyasunaga/BioLinkBERT-large^[8]^. Templates were encoded with the formal Stage 1 maximum of 256 wordpieces. The 1,024-dimensional pooled BioLinkBERT representation was projected by the Stage 1 text adapter into the 256-dimensional AMPHORA condition space. LucaOne and BioLinkBERT encoders were used to generate cached condition features. They were not fine-tuned during Stage 4, where the trainable components were the condition builder, condition projections and output heads, the selected flow route and the target-class head. Genome- and text-condition hit rates were 100% in the final Stage 4 manifests.

### Peptide representation and latent generation

smORFLLM was used as the peptide representation backbone throughout AMPHORA. The encoder was a 30-layer transformer with hidden dimension 640, 20 attention heads, multilayer-perceptron expansion ratio 4.0 and dropout 0.05. The amino-acid vocabulary contained 32 tokens, including padding, beginning-of-sequence, end-of-sequence, unknown and mask tokens. Sparse biophysical fusion used 17 global peptide descriptors and 5 residue-level descriptors. The global descriptors were net charge at pH 7.0, mean Kyte–Doolittle hydropathy, isoelectric point, aromaticity, instability index, Boman-style binding index, alpha-helical hydrophobic moment, charge density, hydrophobicity distribution, helix propensity fraction, sheet propensity fraction, coil propensity fraction, disorder mean, disorder fraction, disorder maximum, mean transmembrane-proxy probability and transmembrane-proxy segment count. The residue-level descriptors were binary flags for hydrophobic, positively charged, flexible, polar and disorder-prone residues. These sequence-derived priors follow recent AMP prediction and design workflows^[5;11;12]^.

smORFLLM was pretrained with masked language modelling and auxiliary biophysical objectives. The reported checkpoint used 150,000 optimization steps, global batch size 8,192 and bf16 distributed training. Frozen-transfer checks used PepBERT9-style peptide classification tasks^[13]^. Encoder weights were fixed, residue states were mean-pooled and a linear probe was trained for each of the nine tasks using shared task manifests and seeds {42, 43, 44}. This comparison assessed representation accessibility for the selected smORFLLM checkpoint and reduced-budget architecture ablations. It was not used as a downstream AMP-generation endpoint.

Stage 1 aligned peptide, genome and text modalities to a shared peptide-conditioned space using adapter and projection modules. The peptide backbone was LoRA-tuned^[14]^ with rank 8, alpha 16 and dropout 0. Peptide, genome and text vectors were projected into 256 dimensions, ℓ_2_-normalized and aligned with positive masks based on instance identity, shared species/taxon/target class and shared task or target class. The full contrastive formulation is given in Supplementary Methods.

Stage 2 converted smORFLLM hidden states into the 256-dimensional latent sequence used by the flow models. Training used an unconditional cluster50 AMP corpus with 1,000,000 sampled training sequences and 10,000 validation sequences. The peptide backbone was frozen. A projection module generated latent tokens, a decoder reconstructed amino-acid strings and a preservation head reconstructed detached smORFLLM hidden states.

Stage 3 trained unconditional flow-matching models over the Stage 2 latent space^[15]^. The training corpus was a predicted-AMP union set derived from the upstream smORFLLM pretraining sequence pool, screened with stored PepNet and AMPlify AMP-prediction calls, together with AVP-IFT-new calls from an in-house AVP-IFT implementation retrained on ESCAPE multilabel AMP labels^[10;12;16;17]^. The de-duplicated and cluster50 split contained 9,388,496 training sequences and 47,180 validation sequences. Predictor-derived labels were used only for corpus construction. Four retained flow families were compared: sequence-only, global-structure latent, joint sequence–structure and pair-support dual-stream models. The pair-support dual-stream model was selected because it gave the best balance of decoding quality, AMP-like neighbourhood support and predicted structure features.

Before conditional training, the selected pair-support model was adapted on a real-AMP split collected from APD, CAMP_R4_, DBAASP, DRAMP and dbAMP^[18–22]^. Merged sequences were normalized, de-duplicated and split after MMseqs2 clustering at 40% identity^[9]^, yielding 40,760 training peptides, 5,094 validation peptides and 5,094 held-out peptides. The low-rank route tuned the last 12 flow layers and selected output, normalization, velocity, decoder, preservation and structure heads. The peptide backbone and input/projector modules remained frozen.

### Conditional generation

Stage 4 introduced target context while retaining the selected sequence–structure generative prior. The input dataset was the ESCAPE-derived conditional corpus. Final conditional runs used rebuilt aligned genome and text embeddings from the Stage 1 alignment model and a separate target-class slot. The training and validation manifests had 100% genome-and text-condition hit rates, exact train–validation peptide-sequence overlap 0 and exact train–validation peptide–target-class-pair overlap 0.

Target context was represented by target class, genome condition and text condition. AMPHORA-OS used all three orthogonal condition slots. Partial conditions used target class only, target class plus genome or target class plus text. Classifier-free condition dropout removed condition slots with probability 0.1 during training^[23]^. In the final AMPHORA-OS run, the target-class slot used an additional dropout probability of 0.6. Genome-slot taxon-context fusion and target-class-fusion shortcuts were disabled. Alignment heads and the peptide decoder were fixed. The condition builder, condition projections, output heads, selected flow route and target-class head were trainable.

The final AMPHORA-OS configuration used the real-AMP-LoRA-derived model-only initialization lineage, learning rate 2 × 10^−4^, cosine decay, 1,000 warm-up steps, batch size 32, bf16 autocast, gradient clipping 0.5, checkpoint interval 500 steps and top-10 checkpoint retention. The target-class auxiliary loss weight was 0.25. Validation target-class balanced accuracy selected the checkpoint used for final evaluation, preventing the majority bacterial class from dominating checkpoint selection.

Formal four-class evaluations sampled up to 250 validation conditions per target class and four generated peptides per condition, producing 2,940 generated sequences because the validation split contained only 215 viral and 20 parasite conditions. AMPHORA-OS used 16 sampling steps and guidance scale 2.5 in the final condition-control analyses. Temperature was 1.0 unless otherwise stated. The sampler initialized latent variables from Gaussian noise, integrated from *t* = 1 to *t* = 0 and decoded the final latent greedily. No post-hoc reranking was applied in raw target-class assignment evaluations unless explicitly stated.

### Generator benchmarks and condition controls

The fixed-25aa benchmark compared generators under a common constant-length protocol. AMPHORA pair-support fixed-25aa and an AMP-Diffusion-style fixed-25aa model were trained from scratch on the same 40%-identity-clustered real-AMP split^[24]^. Public HydrAMP, PepCVAE and AMPGAN sequence sets were evaluated with the same protocol^[5;25;26]^, and a real-AMP reference row served as the training-reference anchor. Trainable rows generated peptides at target length 25 with five seeds, 42–46, and 1,000 sequences per seed. Public rows used five non-overlapping subsets of 1,000 sequences after applying the maximum-length-25 filter.

All fixed-length rows were evaluated with the same sequence, function and distribution metrics. Sequence summaries included length, pairwise edit distance, unique 3-mer count, global uniqueness and minimum edit distance to the real-AMP training set. FPD-ESM2 measured embedding-distribution distance to reference peptides^[27]^. MMD used an RBF kernel with median-distance bandwidth. PRDC precision and recall used *k* = 5 nearest-neighbour radii^[28]^. Function summaries used HydrAMP^[5]^, Macrel^[29]^ and APEX outputs.

Conditional assignment was measured with a k-nearest-neighbour target-class classifier in ESM2-t33-650M peptide embedding space^[27]^. Generated and reference peptides were embedded by mean pooling sequence embeddings. Neighbours were ranked by cosine distance, with *k* = min(5, *n*_reference_). Macro F1 was the unweighted mean of class F1 values, and balanced accuracy was the unweighted mean of class recalls. Parasite-associated records were retained in the prespecified four-class summaries. Because the parasite validation subset contained only 20 conditions and 80 generated peptides, parasite-specific recall is reported for transparency but was not used as an independent class-level conclusion. These metrics are embedding-space target-class assignment proxies, not biological activity measurements.

Condition-control analyses used matched, global-shuffled and within-target-class-shuffled genome/text settings. The matched setting used aligned target-class, genome and text tensors. The global-shuffled setting randomly reassigned genome/text tensors across the evaluation condition pool. The within-target-class-shuffled setting shuffled genome/text tensors only among conditions with the same target class. Partial-condition analyses evaluated target class alone, target class plus genome and target class plus text. Same-noise counterfactual analysis fixed target class, initial latent noise, solver settings and sampling seed while swapping one condition channel between same-target-class condition pairs. It used 26 condition pairs and five seeds per pair, yielding 130 paired rows.

### Species-level and APEX analyses

The 12-species species-aware enrichment benchmark evaluated fresh matched generation from the final AMPHORA-OS checkpoint. Twelve target species were evaluated with three seeds and 1,000 generated peptides per species/seed. The canonical name *Nakaseomyces glabratus* was used for the species formerly referred to as *Candida glabrata*. Reference peptides were drawn from the combined train and validation conditional manifests for the same species after sequence de-duplication within species. Each species contributed 1,000 reference entries; species with fewer than 1,000 unique reference sequences were sampled with replacement.

Reference peptides were embedded with the Stage 4 clean latent readout. The preferred multi-prototype readout clustered each species into up to 10 prototypes with MiniBatchKMeans before assigning soft probabilities to generated peptides. Diagonal enrichment was the matched assignment probability to the target species minus the mean off-target assignment probability. Specificity ratio was the matched assignment probability divided by the mean off-target probability. The multi-prototype readout was used as the conservative main-text readout because it reduces sensitivity to a single centroid for heterogeneous AMP reference sets.

APEX scoring used APEX 1.1, the updated bacterial-strain-specific antimicrobial activity predictor derived from the APEX framework^[30;31]^. We used the released APEX pathogen implementation and all eight pretrained ensemble checkpoints distributed with the model. Predictions from the eight base learners were averaged. APEX 1.1 was trained by its authors on peptide sequences with pathogen-specific MIC measurements and public AMP/non-AMP data, including 15,718 MIC values from 1,642 peptides across 11 pathogenic strains. We used these released checkpoints only as an external post-hoc evaluator and did not fine-tune APEX on AMPHORA outputs.

The seven-species APEX benchmark sampled 1,000 peptides for each species, seed and condition setting across seven bacterial species, three seeds and two condition modes: matched and target class only. The reported comparison generated 42,000 peptides before APEX filtering. APEX returned 11 strain-level MIC-style prediction columns. These columns were grouped into seven species-level target sets: one output each for *Acinetobacter baumannii*, *Klebsiella pneumoniae*, *Enterococcus faecalis* and *Enterococcus faecium*; three outputs for *Escherichia coli*; two outputs for *Pseudomonas aeruginosa*; and two outputs for *Staphylococcus aureus*. The species-level predicted score was the arithmetic mean of log_10_ MIC across the strain columns assigned to that species. Peptides longer than 50 residues were skipped by APEX, leaving 41,180 scored sequences across matched and target-class-only rows. APEX scores are external predicted-activity estimates, not wet-lab MIC measurements.

### Novelty, structure and statistics

Validity was the fraction of generated strings containing only canonical amino-acid tokens. Uniqueness was the fraction of non-duplicate generated sequences. Exact novelty was the fraction of generated sequences absent from the training set. The identity< 70% statistic used nearest-training n-gram retrieval followed by edit-distance refinement. Additional split-structure analyses compared training and validation partitions across peptide sequence, species name, taxon identifiers, genus, genome cache key and text cache key.

Structures for generated peptides were predicted with OmegaFold^[32]^. Generated structures were compared with reference structures using Foldseek^[33]^. Structure summaries included pLDDT, secondary-structure fractions, radius of gyration, contact density, Foldseek hit rate, top-hit TM-score and novel-fold fraction under the stated novelty rule. Fig. 3e used the final AMPHORA-OS Stage 4 structure panel, with 1,536 generated peptides and 1,259 reference peptides from the matched full-condition evaluation set. The Stage 3 no-condition versus Stage 4 AMPHORA-OS comparison was treated separately as a lineage-level structure check rather than as the source for Fig. 3e. Because short peptides can be flexible, structure prediction and Foldseek novelty were treated as computational screening readouts.

Condition contrasts in Fig. 2e were computed directly from the final Stage 4 partial-condition and shuffled-condition point estimates reported in Supplementary Tables 6 and 11. Same-noise edit-distance intervals were computed from the paired counterfactual rows, and species-level summaries were aggregated across the prespecified seeds. These uncertainty summaries describe evaluation-sample variation for computational readouts and do not replace repeated training seeds.

## Data availability

The peptide training manifests, generated sequence tables, evaluation splits and source data underlying the main figures and supplementary tables will be deposited in a public repository. Sequence resources integrated from public databases were obtained from APD3, CAMPR4, DBAASP, DRAMP and dbAMP under their respective access conditions. Where redistribution of third-party sequence resources is restricted, accession lists, processed manifests or scripts for reproducing the processed datasets will be provided instead of redistributing the original database records. Source data supporting the figures and supplementary analyses will be provided with this preprint where permitted.

## Code availability

Custom code for model training, conditional generation, APEX-based post-hoc evaluation and downstream analyses will be released in the same public repository. Until repository release, code central to reproducing the main computational analyses is available from the corresponding authors upon reasonable request.

