## Supplementary Information for "Pathogen context reshapes antimicrobial peptide generation"

#### Supplementary Methods

##### Peptide datasets and preprocessing

AMPHORA was trained through four stages with separate objectives. Stage 1 aligned peptide, genome and text representations on a curated conditional AMP corpus. Stage 2 compressed peptide hidden states into a decodable 256-dimensional latent sequence. Stage 3 trained unconditional latent-flow generators and selected the sequence–structure backbone used for conditional modelling. Stage 4 introduced target-class, genome and text conditions through orthogonal condition slots and evaluated matched, partial and shuffled condition settings. These stages were separately optimized but lineage-linked: later stages reused selected backbones, latent spaces or initialization checkpoints from earlier stages as specified below. Trainable and frozen components were recorded for each stage to distinguish representation learning, latent compression, unconditional generation and conditional adaptation. Unless otherwise stated, peptide sequences were represented with the 20 standard amino acids and were capped at 128 residues.

The peptide representation model was pretrained on a short-peptide/smORF corpus rather than on AMP activity labels. Preprocessing started from a representative FASTA containing 390,709,416 sequences assembled from public short-protein/smORF resources and internal smORF collections. MMseqs2 cluster-consistent splitting at 50% sequence identity produced 386,797,786 training rows, 1,951,071 validation rows and 1,960,384 test rows in the shard manifest[1]. Length-aware sampling retained all sequences of length at most 50, retained sequences of length 51–90 with probability 0.5, and retained longer sequences with probability 0.2. The maximum sequence length for model input was 128 residues.

This corpus was not AMP-specific, and smORFLLM pretraining did not use downstream AMP activity labels. Because public sequence sources may contain AMP-like sequences, downstream AMP-specific train–test overlap was evaluated separately with exact-match and low-identity screens.

The conditional corpus used for peptide–pathogen alignment was derived from the AMP-positive subset of ESCAPE, a standardized multilabel AMP benchmark integrating more than 80,000 peptides from 27 validated repositories and annotating antibacterial, antifungal, antiviral and antiparasitic functions[2]. The ESCAPE-derived AMP-positive source table contained 21,409 unique AMP sequences. Peptide–target pathogen conditions were constructed by combining dbAMP and DRAMP target annotations with ESCAPE category labels, followed by manual checks for pathogen/category consistency. Exact dbAMP or DRAMP matches were used as target-annotation evidence. Records supported only by ESCAPE category fallback labels were retained as target-class or category-level conditions rather than as validated species-specific activity evidence.

The formal split contained 93,834 training rows and 4,910 validation rows, corresponding to 20,313 and 1,070 unique peptides, respectively. Exact train–validation peptide-sequence overlap was zero. The training set contained 73,500 bacterial, 13,745 fungal, 5,999 viral and 590 parasite rows. The validation set contained 3,842 bacterial, 833 fungal, 215 viral and 20 parasite rows. These counts define the long-tailed structure of the conditional corpus and were retained in the prespecified four-class analyses.

#### Pathogen-context annotation

Manually checked conditions were expanded into peptide–pathogen–genome–text records. Each retained row was assigned a canonical species or taxon identifier, a representative genome cache key and a standardized strain-description text template through a manually curated pathogen mapping table. For rows without validated species-specific activity evidence, the target class was retained as category-level evidence and the genome/text condition was assigned through the curated mapping. Each row contained a peptide sequence, taxonomy identifier, target class, species identifier, genome cache key, text cache key and text template. Task mask, route identifier and activity-bin metadata were retained as provenance and stratification fields.

#### Genome-context processing

Genome conditions were cached before conditional generation and paired to peptide–pathogen records through the curated genome cache key. For each cache key, the representative genome FASTA was encoded from sampled nucleotide windows. Contigs were split into windows of up to 4,096 nt with a 2,048-nt stride, retaining up to 64 windows through uniform, high-complexity and deterministic random sampling. Raw genome embeddings were produced with the LucaOne gene encoder checkpoint `LucaGroup/LucaOne-gene-step36.8M`[3], using `seq_type="gene"` and mean pooling across sampled windows. The raw LucaOne genome representation was 2,560-dimensional. The exported Stage 1 aligned genome adapter projected this representation to the 256-dimensional AMPHORA condition space and retained aligned window embeddings when available. Stage 1 used cached raw genome features together with peptide and text features to learn modality projections into peptide space rather than fine-tuning the full LucaOne encoder. In the final Stage 4 runs, genome embeddings were fixed cached inputs; trainable components included the condition builder, condition projections and output heads, the selected flow route and the target-class head. Genome-condition hit rate was 100% in the final Stage 4 training and validation manifests.

#### Text-context processing

Text conditions were cached before conditional generation and paired to peptide–pathogen records through the curated text cache key and standardized strain-description template. Text embeddings were produced from whitespace-normalized strain-description templates with the BioLinkBERT-large checkpoint `michiyasunaga/BioLinkBERT-large`[4], using the formal Stage 1 maximum of 256 wordpieces. The raw BioLinkBERT pooled representation was 1,024-dimensional and was stored as a single text token. The exported Stage 1 aligned text adapter projected this representation to the 256-dimensional AMPHORA condition space. Stage 1 used cached raw text features together with peptide and genome features to learn modality projections into peptide space rather than fine-tuning the full BioLinkBERT encoder. In the final Stage 4 runs, text embeddings were fixed cached inputs; trainable components included the same condition-builder and flow-route modules described above. Text-condition hit rate was 100% in the final Stage 4 training and validation manifests.

#### Short-peptide representation model

smORFLLM was used as the peptide representation backbone throughout AMPHORA. The encoder was a 30-layer transformer with hidden dimension 640, 20 attention heads, multilayer-perceptron expansion ratio 4.0 and dropout 0.05. The amino-acid vocabulary contained 32 tokens, including padding, beginning-of-sequence, end-of-sequence, unknown and mask tokens. Sparse biophysical fusion used 17 global peptide descriptors and 5 residue-level descriptors. Global and residue-level priors were injected through explicit fusion layers at layers 20 and 30, with fusion repeated every four layers in the configured sparse-fusion schedule. The residual variant used block attention residuals with block size 5.

Global biophysical descriptors were deterministic sequence-derived priors rather than activity labels. Before descriptor calculation, sequences were uppercased and filtered to the 20 standard amino acids. The 17-dimensional fingerprint contained, in order, net charge at pH 7.0, mean Kyte–Doolittle hydropathy, isoelectric point, aromaticity, instability index, Boman-style binding index, alpha-helical hydrophobic moment, charge density, hydrophobicity distribution, helix propensity fraction, sheet propensity fraction, coil propensity fraction, disorder mean, disorder fraction, disorder maximum, mean transmembrane-proxy probability and transmembrane-proxy segment count. These descriptor families follow recent AMP prediction and design workflows that use composition, charge, hydrophobicity, amphipathicity and related physicochemical features as sequence-derived inputs[5–7]. Aromaticity and instability index were computed with the Biopython `ProteinAnalysis` implementation when available. Hydropathy-derived quantities used the Kyte–Doolittle scale. Hydrophobic moment used a 100-degree rotation for an ideal alpha helix. Secondary-structure fractions used Chou–Fasman-style propensities. The binding index was implemented as a Boman-style residue-potential average. Transmembrane quantities were lightweight hydropathy-window proxies. The five residue-level descriptors were rebuilt online from the token sequence as binary flags for hydrophobic residues (A, I, L, M, F, W, V and Y), positively charged residues (K, R and H), flexible residues (G, S, P, D, N and Q), polar residues (S, T, N, Q, D, E, K, R and H) and disorder-prone residues (P, G, S, Q, E, K and A).

#### smORFLLM pretraining

smORFLLM was pretrained with masked language modelling and auxiliary biophysical objectives. Tokens were selected for masking with probability 0.15. Selected tokens were replaced by the mask token with probability 0.8, by a random token with probability 0.1 and left unchanged otherwise. The training objective was

$$\mathcal{L}_{\text{smORF}} = \mathcal{L}_{\text{MLM}} + 0.01\,\mathcal{L}_{\text{bio\_reg}} + 0.01\,\mathcal{L}_{\text{bio\_cls}}, \tag{1}$$

where $\mathcal{L}_{\text{MLM}}$ is token cross-entropy on masked positions, $\mathcal{L}_{\text{bio\_reg}}$ is the auxiliary regression loss for biophysical descriptors and $\mathcal{L}_{\text{bio\_cls}}$ is the auxiliary biophysical classification loss.

The reported checkpoint used 150,000 optimization steps, global batch size 8,192 and gradient accumulation 2. The optimizer was AdamW with learning rate $1.2 \times 10^{-4}$, minimum learning rate $1.2 \times 10^{-5}$, 10,000 warm-up steps, weight decay 0.1 and $\beta = (0.9, 0.95)$. Gradients were clipped at 1.0. Distributed training used bf16 precision.

Frozen-transfer checks used PepBERT9-style peptide classification tasks[8]. Encoder weights were fixed, residue states were mean-pooled and a linear probe was trained for each of the nine tasks. The same task manifest, probe family, seeds $\{42, 43, 44\}$, probe batch size 512 and short-peptide cap of 50 residues were used for all compared models. This comparison was used to assess representation accessibility for the selected smORFLLM checkpoint and reduced-budget architecture ablations, not as a downstream AMP-generation endpoint.

#### Peptide–genome–text alignment

Stage 1 aligned peptide, genome and text modalities to a shared peptide-conditioned space using adapter and projection modules. The peptide backbone was the smORF150M-derived encoder with LoRA tuning[9]. LoRA rank was 8, alpha was 16 and dropout was 0. Training used learning rate $2 \times 10^{-4}$, batch size 32, gradient accumulation 1, 3 epochs, checkpoint interval 200 steps and 32 data-loader workers. The alignment objective was active; sequence, flow, structure, self-flow and preservation losses were disabled.

For a minibatch of size $B$, each available modality $m$ produced one 256-dimensional projected vector per item. Peptide vectors were obtained by attention-mask mean pooling of peptide hidden states followed by a peptide projection head. Genome and text vectors were pooled from cached tokens and passed through modality-specific projection heads. Target class, species and task metadata defined positive masks. All vectors were $\ell_2$-normalized before similarity calculation.

For modalities $m$ and $n$, a positive mask $M$ defined positive cross-modal pairs and

$$q_{ij}^{m,n} = \exp((a_i^m)^\top a_j^n/\tau), \tag{2}$$

with $\tau = 0.07$. The directed log-sum-exp positive loss was

$$\ell_{m\to n}(M) = -\frac{1}{|\mathcal{I}_M|}\sum_{i\in\mathcal{I}_M} \log\left(\frac{\sum_j M_{ij} q_{ij}^{m,n}}{\sum_j R_{ij} q_{ij}^{m,n}}\right), \tag{3}$$

where $R$ is the valid-pair mask after missing-modality filtering and $\mathcal{I}_M$ contains rows with at least one positive pair. The bidirectional term was

$$\ell_{m,n}(M) = \frac{1}{2}\{\ell_{m\to n}(M) + \ell_{n\to m}(M^\top)\}. \tag{4}$$

Three positive masks were used: instance identity, shared species/taxon/target-class group and shared task or target class. The Stage 1 alignment loss was averaged across available modality pairs and positive-mask levels.

#### Latent peptide compression and decoding

For a peptide sequence $x = (x_1, \ldots, x_L)$, the smORFLLM encoder produced residue-level hidden states $H_\theta(x) \in \mathbb{R}^{L\times 640}$. A projection module mapped these states into the latent sequence used by the flow models,

$$z_0 = P_\psi(H_\theta(x)) \in \mathbb{R}^{L\times 256}. \tag{5}$$

Stage 2 converted smORFLLM hidden states into the 256-dimensional latent sequence used by the flow models. Training used an unconditional cluster50 AMP corpus rather than the conditional manifest. The formal Stage 2 training set contained 1,000,000 sampled sequences and the validation set contained 10,000 sequences. Precomputed peptide hidden caches were produced by the Stage 1 peptide backbone after merging the peptide LoRA weights. Genome, text and OmegaFold-derived structure caches were disabled.

The projector $P_\psi$ was a shallow residual projection network with input layer normalization, a two-layer MLP with hidden dimension 512, a residual linear branch, output layer normalization and an optional depthwise convolution with kernel size 3. The decoder $D_\phi$ reconstructed amino-acid tokens from the latent sequence. The preservation head $R_\rho$ reconstructed detached smORFLLM hidden states. For attention mask $m_i$, the sequence and preservation losses were

$$\begin{aligned} \mathcal{L}_{\text{seq}} &= \text{CE}(D_\phi(z_0), x), \\ \mathcal{L}_{\text{preserve}} &= \left(\sum_i m_i\right)^{-1} \sum_i m_i \|R_\rho(z_{0,i}) - H_{\theta,i}(x)\|_2^2. \end{aligned} \tag{6}$$

A whitening regularizer was applied to valid latent tokens. If $\mu$ and $\Sigma$ denote the empirical mean and covariance of normalized latent tokens in a minibatch, the regularizer was

$$\mathcal{L}_{\text{white}} = \|\mu\|_2^2 + \|\text{diag}(\Sigma) - \mathbf{1}\|_2^2 + \lambda_{\text{cov}}\|\Sigma - \text{diag}(\Sigma)\|_F^2. \tag{7}$$

The Stage 2 objective was

$$\mathcal{L}_{\text{stage2}} = \mathcal{L}_{\text{seq}} + 0.25\,\mathcal{L}_{\text{preserve}} + 0.02\,\mathcal{L}_{\text{white}}. \tag{8}$$

The peptide backbone was frozen. Training used 4 NPUs, total batch size 1,536, per-device batch size 384, validation batch size 1,536, learning rate $10^{-3}$, 3 epochs, gradient accumulation 1, checkpoint interval 500 steps and log interval 50 steps.

#### Unconditional latent-flow training

Stage 3 trained unconditional flow-matching models over the Stage 2 latent space[10]. The training corpus was a predicted-AMP union set derived from the upstream smORFLLM pretraining sequence pool. The short-peptide/smORF collection was screened with stored PepNet and AMPlify AMP-prediction call sets, together with AVP-IFT-new calls from an in-house implementation of the AVP-IFT architecture retrained on ESCAPE multilabel AMP labels[2;6;11;12]. A sequence was retained when at least one call set classified it as AMP-positive. AVP-IFT-new therefore denotes the ESCAPE-retrained internal checkpoint rather than an unmodified external antiviral-peptide model. This predictor-union set was de-duplicated and clustered at 50% sequence identity with MMseqs2 before train–validation splitting[1], yielding 9,388,496 training sequences and 47,180 validation sequences. Predictor-derived labels were used only for corpus construction; Stage 3 optimization used the flow, decoding and structure objectives described below.

For a clean latent $z_0$, Gaussian noise $\epsilon \sim \mathcal{N}(0, I)$ and time $t \sim \mathcal{U}(0, 1)$, the noised latent was

$$z_t = (1 - t)z_0 + t\epsilon. \tag{9}$$

The vector field $v_\omega$ was trained to predict the data-to-noise velocity $\epsilon - z_0$:

$$\mathcal{L}_{\text{flow}} = \left(\sum_i m_i\right)^{-1} \sum_i m_i \|v_\omega(z_{t,i}, t) - (\epsilon_i - z_{0,i})\|_2^2. \tag{10}$$

At generation time, sampling was initialized from Gaussian noise at $t = 1$ and integrated backward to $t = 0$. Heun integration was used unless otherwise specified. In conditional sampling, classifier-free guidance combined unconditional and conditional vector fields[13] as

$$v_{\text{cfg}} = v_{\text{uncond}} + s\,(v_{\text{cond}} - v_{\text{uncond}}), \tag{11}$$

where $s$ is the guidance scale.

Four retained Stage 3 families were evaluated. The sequence-only latent flow served as the minimal generative backbone. The global-structure latent flow added sequence-level structural information without residue-pair supervision. The joint sequence–structure latent flow trained sequence and structure latents together. The pair-support dual-stream latent flow maintained separate sequence and structure streams, fused them from the first layer and added pair-support supervision derived from OmegaFold pair-bias features. All four families were trained on the same Stage 2 peptide latent cache, used the same cluster50 train–validation split and excluded target-class, genome and text conditions.

Checkpoints were selected by validation pseudo-perplexity and evaluated with a common protocol. For each retained model, 10,000 generated peptides were compared with 10,000 reference peptides sampled from the same evaluation scaffold. The reported checkpoint steps were 17,000 for the sequence-only flow, 13,500 for the global-structure flow, 22,000 for the joint sequence–structure flow and 14,500 for the pair-support dual-stream flow. Validity, uniqueness and exact train novelty were computed before distributional evaluation; all four retained models reached 1.000 for these checks in the Stage 3 evaluation.

The selected pair-support backbone used dual sequence and structure latent streams. It predicted sequence and structure-latent velocities, fused the streams from the first layer, applied token dropout 0.05, used hidden dimension 768 and included a pair-support branch. The structure stream used the same interpolation for an Omega-derived structure latent $s_0$, with $\epsilon^s \sim \mathcal{N}(0, I)$ and $s_t = (1-t)s_0 + t\epsilon^s$. The structure-flow loss was

$$\mathcal{L}_{\text{flow\_struct}} = \left(\sum_i m_i\right)^{-1} \sum_i m_i \|v^s_\omega(s_{t,i}, t) - (\epsilon^s_i - s_{0,i})\|_2^2. \tag{12}$$

Auxiliary structure supervision used a smooth-$L_1$ residue-structure loss and a pair-support binary cross-entropy loss. Pair-support targets were defined by non-zero Omega-derived pair-bias entries, diagonal positions were masked and the positive class weight was 4.0. The selected pair-support objective used weights 1.0 for sequence flow, 1.0 for structure flow, 0.5 for sequence decoding, 0.25 for hidden-state preservation, 0.25 for single-residue structure supervision and 0.05 for pair-support supervision.

All Stage 3 variants used learning rate $10^{-3}$, cosine decay, 1,000 warm-up steps, minimum learning-rate ratio 0.1, gradient clipping 0.5, 3 epochs, checkpoint interval 500 steps and pseudo-perplexity as the best-checkpoint metric. Simpler variants used per-device batch size 320 across 4 devices, giving total batch size 1,280. Dual-stream variants used per-device batch size 160 with gradient accumulation 2, also giving effective batch size 1,280.

#### Real-AMP adaptation before conditional training

After backbone selection, the pair-support dual-stream model was adapted on a real-AMP split by low-rank fine-tuning before Stage 4. Real-AMP sequences were collected from APD, $\text{CAMP}_{\text{R4}}$, DBAASP, DRAMP and dbAMP resources[14–18]. Merged sequences were normalized, de-duplicated and split after MMseqs2 clustering at 40% identity[1]. The split contained 40,760 training peptides, 5,094 validation peptides and 5,094 held-out peptides. The held-out partition was not used in the LoRA training loop and was used only where explicitly stated in downstream reference construction.

The adaptation initialized from the selected pair-support Stage 3 model-only checkpoint. Target-class, genome and text condition caches were disabled. Cached peptide hidden states and OmegaFold-derived structure caches from the real-AMP route were used. The low-rank route tuned the last 12 flow layers with LoRA rank 12, alpha 24 and dropout 0.05. Output projections, normalization layers, velocity heads, the decoder, the preservation head and structure heads were trainable; the peptide backbone and input/projector modules were frozen. Training used learning rate $3 \times 10^{-6}$, 12 epochs, batch size 160, validation batch size 320, gradient accumulation 2, cosine scheduling, 400 warm-up steps and gradient clipping 0.5. The objective was

$$\mathcal{L}_{\text{real-AMP}} = \mathcal{L}_{\text{flow}} + \mathcal{L}_{\text{flow\_struct}} + 0.5\,\mathcal{L}_{\text{seq}} + 0.45\,\mathcal{L}_{\text{preserve}} + 0.25\,\mathcal{L}_{\text{of\_single}} + 0.05\,\mathcal{L}_{\text{of\_pair}}. \quad (13)$$

The resulting real-AMP-adapted model-only checkpoint supplied the source weights for Stage 4 conditional training.

#### Conditional flow generation

Target context was represented by target class $y$, genome condition $g$ and text condition $u$. The condition builder assembled these inputs as

$$c = B_\eta(y, g, u). \quad (14)$$

The four target classes used in the formal analyses were bacterium, fungus, virus and parasite.

Stage 4 introduced pathogen context while retaining the selected sequence–structure generative prior. The input dataset was the ESCAPE-derived conditional corpus described above. Final conditional runs used rebuilt aligned genome and text embeddings from the Stage 1 alignment model and a separate target-class slot. The training and validation manifests had 100% genome-condition and text-condition hit rates, exact train–validation peptide-sequence overlap 0 and exact train–validation peptide–target-class-pair overlap 0.

The Stage 4 weight lineage was defined by checkpoint role. Stage 3 selected the pair-support backbone, which was then adapted on the real-AMP split and exported as the real-AMP-adapted pair-support generator. For conditional training, the real-AMP-LoRA-derived lineage was converted through an intermediate full-condition Stage 4 route into a model-only initialization. Optimizer state was removed and the training step reset before downstream Stage 4 training. The clean full-condition base generator and AMPHORA with orthogonal condition slots (AMPHORA-OS) were trained separately from this initialization lineage; AMPHORA-OS was not continued from the subsequently trained base-generator checkpoint.

The condition builder assembled target-class, genome and text slots as

$$c = B_\eta(y, g, u) = [c_{\mathrm{tc}}, c_{\mathrm{genome}}, c_{\mathrm{text}}]. \tag{15}$$

AMPHORA-OS used all three slots. Partial conditions used target class only, target class plus genome or target class plus text. Classifier-free condition dropout removed condition slots with probability 0.1 during training. In the final AMPHORA-OS run, the target-class slot used an additional dropout probability of 0.6. Genome-slot taxon-context fusion and target-class-fusion shortcuts were disabled. The sequence and structure condition scales were 1.0 and 0.8, respectively. Alignment heads and the peptide decoder were fixed. The condition builder, condition projections and output heads, selected flow route and target-class head were trainable.

The conditional vector field minimized the same latent flow-matching objective conditioned on $c$:

$$\mathcal{L}_{\mathrm{cond\_flow}} = \left(\sum_i m_i\right)^{-1} \sum_i m_i \left\| v_\omega(z_{t,i}, t, c) - (\epsilon_i - z_{0,i}) \right\|_2^2 . \tag{16}$$

The denoised latent was $\hat{z}_0 = z_t - t\, v_\omega(z_t, t, c)$. The target-class head predicted $y$ from pooled $\hat{z}_0$. Conditional sequence and preservation losses were

$$\begin{aligned} \mathcal{L}_{\mathrm{seq}}^{\mathrm{cond}} &= \mathrm{CE}(D_\phi(\hat{z}_0), x), \\ \mathcal{L}_{\mathrm{preserve}}^{\mathrm{cond}} &= \left(\sum_i m_i\right)^{-1} \sum_i m_i \left\| R_\rho(\hat{z}_{0,i}) - H_{\theta,i}(x) \right\|_2^2 . \end{aligned} \tag{17}$$

The Stage 4 objective was

$$\mathcal{L}_{\mathrm{stage4}} = \mathcal{L}_{\mathrm{cflow}} + 0.5\, \mathcal{L}_{\mathrm{seq}}^{\mathrm{cond}} + 0.1\, \mathcal{L}_{\mathrm{preserve}}^{\mathrm{cond}} + 0.25\, \mathcal{L}_{\mathrm{tc}}, \tag{18}$$

where $\mathcal{L}_{\mathrm{tc}}$ was cross-entropy over the four target classes.

The clean full-condition Stage 4 base-generator route was trained on RTX 4090D-class CUDA hardware with batch size 32, bf16 autocast, learning rate $2 \times 10^{-4}$, cosine decay, 1,000 warm-up steps, gradient clipping 0.5, 3 epochs, target-class strength 1.25 and top-10 checkpoint retention. Target-class strength is the scalar multiplier applied to the embedded target-class token before slot normalization and fusion with cached genome and text tokens. The best checkpoint was selected by validation target-class balanced accuracy,

$$\mathrm{BA} = \frac{1}{4} \sum_{k=1}^{4} \frac{\mathrm{TP}_k}{\mathrm{TP}_k + \mathrm{FN}_k}. \tag{19}$$

This metric prevented the majority bacterial class from dominating checkpoint selection.

AMPHORA-OS used the same real-AMP-LoRA-derived model-only initialization lineage as the full-condition route, with training state reset before optimization. The final configuration used a full Stage 4 update with condition target-class strength 1.25, classifier-free condition dropout 0.1[13], target-class-slot dropout 0.6, no length-bucket condition token and zero taxon, genome and text residual weights outside the orthogonal slots. The target-class auxiliary loss weight was 0.25 with ordinary cross-entropy. Training used learning rate $2 \times 10^{-4}$, cosine decay, 1,000 warm-up steps, batch size 32, bf16 autocast, gradient clipping 0.5, checkpoint interval 500 steps and top-10 checkpoint retention. Validation target-class balanced accuracy selected the checkpoint used for final evaluation.

#### Conditional peptide sampling

Formal four-class evaluations sampled up to 250 validation conditions per target class and four generated peptides per condition. This yielded 2,940 generated sequences because the validation split contained only 215 viral and 20 parasite conditions. Reference panels were sampled from the same validation split used for generation, with up to 512 reference peptides per class where available. AMPHORA-OS used 16 sampling steps and guidance scale 2.5 in the final condition-control analyses. Temperature was 1.0 unless otherwise stated. Fixed-25aa analyses used fixed length 25; four-class assignment analyses used the lengths specified by the corresponding evaluation protocol and condition batch.

The sampler initialized $z_1 \sim \mathcal{N}(0, I)$ and, for dual-stream models, $s_1 \sim \mathcal{N}(0, I)$. It then integrated from $t = 1$ to $t = 0$. With Euler integration,

$$z_{t-\Delta t} = z_t - \Delta t \, v_\omega(z_t, t, c). \tag{20}$$

The Heun solver first computed an Euler proposal and then averaged vector fields at the current and proposed next state. The final latent was decoded greedily with the peptide decoder. No post-hoc reranking was applied in raw target-class assignment evaluations unless explicitly stated.

#### Fixed-length AMP generator benchmark

The fixed-25aa benchmark compared generators under a common constant-length protocol. The two trainable models were AMPHORA pair-support fixed-25aa and an AMP-Diffusion-style fixed-25aa model; both were trained from scratch on the same 40%-identity-clustered real-AMP split[19]. Public HydrAMP, PepCVAE and AMPGAN sequence sets were evaluated with the same protocol[7;20;21], and a real-AMP reference row served as the training-reference anchor.

Trainable AMPHORA and AMP-Diffusion-style rows generated peptides at target length 25 with five seeds, 42–46, and 1,000 sequences per seed, producing 5,000 generated sequences per model before metric aggregation. Public HydrAMP, PepCVAE and AMPGAN rows used five non-overlapping subsets of 1,000 sequences from public sequence sets after applying the maximum-length-25 filter. The exact-25aa 40%-identity-clustered real-AMP reference contained 765 pooled train/validation/test sequences. The held-out test-only exact-25aa subset contained 37 sequences, so the pooled exact-25aa reference was used for stable distributional and structure-related comparisons.

All rows were evaluated with the same sequence, function and distribution metrics. Sequence summaries included number of sequences, mean length, length standard deviation, average pairwise edit distance, unique 3-mer count, global unique fraction and average minimum edit distance to the real-AMP training set. FPD-ESM2 was the Fréchet distance between generated and reference ESM2 embeddings[22]; lower values indicate closer embedding-distribution match. MMD used an RBF kernel with median-distance bandwidth. PRDC precision and recall used $k = 5$ nearest-neighbour radii[23]. Function summaries were post-hoc predictor outputs. HydrAMP pAMP and pMIC were the mean predicted AMP probability and low-MIC probability from HydrAMP[7]. Macrel P(AMP) was the mean AMP probability from Macrel[24]. APEX-predicted average MIC was the mean MIC-style prediction averaged over the 11 bacterial strain outputs available in APEX; lower values are better. APEX-predicted MIC $< 100\ \mu$M was the fraction of generated sequences below this threshold.

#### Target-class assignment analysis

Conditional assignment was measured with a k-nearest-neighbour target-class classifier in peptide embedding space. Generated and reference peptides were embedded with ESM2-t33-650M using mean-pooled sequence embeddings[22]. Let $e(x)$ denote the generated peptide embedding and $r_j$ a reference embedding with label $y_j$. Neighbours were ranked by cosine distance, with $k = \min(5, n_{\text{reference}})$. The predicted class was the majority label among the $k$ nearest reference embeddings. Per-class precision, recall and F1 were computed as

$$P_k = \frac{\text{TP}_k}{\text{TP}_k + \text{FP}_k}, \quad R_k = \frac{\text{TP}_k}{\text{TP}_k + \text{FN}_k}, \quad F1_k = \frac{2P_kR_k}{P_k + R_k}. \tag{21}$$

Macro F1 was the unweighted mean of $F1_k$ over the four target classes, and balanced accuracy was the unweighted mean of class recalls. Parasite-associated records were retained in the prespecified four-class summaries. Because the parasite validation subset contained only 20 conditions and 80 generated peptides, parasite-specific recall is reported for transparency but was not used as an independent class-level conclusion. These metrics are embedding-space target-class assignment proxies, not measurements of biological activity.

#### Matched and shuffled condition sets

Formal condition-control analyses used the same ESM2-kNN assignment proxy. Unless otherwise stated, generation used guidance scale 2.5, 16 sampling steps, up to 250 conditions per target class and four samples per condition. All rows are computational class-alignment readouts rather than biological activity assays.

The condition-shuffle analysis evaluated matched, global-shuffled and within-target-class-shuffled settings. The matched setting used aligned target-class, genome and text condition tensors. The global-shuffled setting randomly reassigned genome/text tensors across the evaluation condition pool. The within-target-class-shuffled setting shuffled genome/text tensors only among conditions with the same target class. Matched versus global-shuffled tested condition use beyond an unconditional peptide prior; matched versus within-target-class-shuffled tested fine-grained genome/text signal beyond coarse target class.

Partial-condition analyses evaluated target class alone, target class plus genome and target class plus text. Matched-versus-partial comparisons tested whether genome and text slots added signal beyond the target-class label. Matched-versus-shuffled comparisons tested whether the additional signal survived condition perturbation.

Same-noise counterfactual analysis fixed target class, initial latent noise, solver settings and sampling seed while swapping one condition channel between same-target-class condition pairs. The analysis used 26 same-target-class condition pairs and five seeds per pair, yielding 130 paired rows. Four perturbations were evaluated: genome-only swap, full-condition genome swap with text fixed, text-only swap and full-condition text swap with genome fixed. Outcomes were decoded-sequence change, edit distance between paired decoded sequences and relative $L_2$ change in the model velocity field.

Same-noise directionality analysis used the same paired generations to test whether a swap moved the generated peptide embedding toward the donor-species prototype. For each perturbation, the change in similarity to the donor prototype was computed after the swap, and 95% confidence intervals were obtained by bootstrap resampling over paired rows. This analysis was treated as a stronger directional test than sequence change alone.

#### Species-level enrichment analysis

The 12-species species-aware enrichment benchmark evaluated fresh matched generation from the final AMPHORA-OS checkpoint. Twelve target species were evaluated with three seeds and 1,000 generated peptides per species/seed. The canonical name *Nakaseomyces glabratus* was used for the species formerly referred to as *Candida glabrata*. Reference peptides were drawn from the combined train and validation conditional manifests for the same species after sequence de-duplication within species. Each species contributed 1,000 reference entries; species with fewer than 1,000 unique reference sequences were sampled with replacement. Because several fungal species had fewer than 1,000 unique reference sequences, the benchmark is interpreted as a balanced computational readout rather than as independent activity validation.

Reference peptides were embedded with the Stage 4 clean latent readout. A single-centroid readout represented each species by one centroid. The preferred multi-prototype readout clustered each species into up to 10 prototypes with MiniBatchKMeans before assigning soft probabilities to generated peptides. For target species $i$, diagonal enrichment was the matched assignment probability to $i$ minus the mean off-target assignment probability. Specificity ratio was the matched assignment probability divided by the mean off-target probability. The multi-prototype readout was used as the conservative main-text readout because it reduces sensitivity to a single centroid for heterogeneous AMP reference sets.

#### Post-hoc bacterial prioritization with APEX

APEX scoring used APEX 1.1, the updated bacterial-strain-specific antimicrobial activity predictor derived from the APEX framework[25;26]. We used the released APEX pathogen implementation and all eight pretrained ensemble checkpoints distributed with the model; predictions from the eight base learners were averaged. APEX 1.1 was trained by its authors on peptide sequences with pathogen-specific MIC measurements and public AMP/non-AMP data, including 15,718 MIC values from 1,642 peptides across 11 pathogenic strains. We used these released checkpoints only as an external post-hoc evaluator and did not fine-tune APEX on AMPHORA outputs.

The seven-species APEX benchmark used the final AMPHORA-OS checkpoint and sampled 1,000 peptides for each species, seed and condition setting across seven bacterial species, three seeds and two condition modes: matched and target class only. The reported comparison generated 42,000 peptides before APEX filtering. APEX returned 11 strain-level MIC-style prediction columns. These columns were grouped into seven species-level target sets: one output each for *Acinetobacter baumannii*, *Klebsiella pneumoniae*, *Enterococcus faecalis* and *Enterococcus faecium*; three outputs for *Escherichia coli*; two outputs for *Pseudomonas aeruginosa*; and two outputs for *Staphylococcus aureus*. For a generated peptide assigned to species $s$, the species-level predicted score was the arithmetic mean of $\log_{10}$ MIC across the APEX strain columns assigned to $s$. Displayed MIC-scale values were obtained by exponentiating this mean log score. Peptides longer than 50 residues were skipped by the APEX evaluator, leaving 41,180 scored sequences across matched and target-class-only rows.

The primary matched-pool utility readout was the fraction of generated peptides with species-level APEX-predicted MIC below 100 $\mu$M. The conditioning-gain readout was matched minus target-class-only mean predicted $\log_{10}$ MIC; negative values indicate lower predicted MIC for matched generation. APEX scores are post-hoc computational prioritization readouts, not wet-lab MIC measurements.

#### Controlled genome and text swaps

Condition-influence maps measured how much each condition slot changed the model velocity field when masked from the full-condition input. Target-class, genome and text slot masks were evaluated across flow times $t \in \{0.05, 0.20, 0.40, 0.60, 0.80, 0.95\}$, transformer layers, 32 conditions and four seeds per condition. For sequence and structure streams separately, the slot influence share was the slot-specific perturbation magnitude divided by the sum across target-class, genome and text slots at the same time/layer. Active-layer summaries excluded layers before condition injection, where all slot shares were zero. Time-trend summaries averaged over all layers to show trajectory-level effects.

#### Novelty and memorization analysis

Validity was defined as the fraction of generated strings containing only canonical amino-acid tokens under the sequence parser. Uniqueness was the fraction of non-duplicate generated sequences. Exact novelty was the fraction of generated sequences absent from the training set. The identity$< 70\%$ statistic used a nearest-training screen based on n-gram retrieval followed by edit-distance refinement. For a generated sequence $x$, the reported nearest-training identity was the maximum identity identified by this screen over the indexed training set.

Additional deterministic quality-control analyses evaluated sequence novelty, split structure, condition perturbation and benchmark provenance. These analyses used the clean Stage 2 train/validation manifests, AMPHORA-OS generated sequence tables, final evaluation summaries and the fixed-25aa benchmark table. Split-structure analysis compared training and validation partitions across exact peptide sequence, canonical species name, canonical species taxon identifier, target taxon identifier, genus, genome cache key and text cache key. For each unit $u$, we computed unique training values, unique validation values, overlapping values and the fraction of validation rows whose unit value appeared in training. This analysis distinguished exact peptide non-overlap from organism-, taxonomy- and condition-cache overlap.

#### Predicted structural plausibility analysis

Structures for generated peptides were predicted with OmegaFold[27]. Generated structures were compared with reference structures using Foldseek[28]. Structure-panel summaries included mean and median pLDDT, fractions of residues and sequences above pLDDT thresholds, predicted secondary-structure fractions, radius of gyration, contact density, fraction of generated queries with a Foldseek hit, top-hit TM-score and novel-fold fraction under the stated novelty rule. TM-score is the template-modelling structural-similarity score for the best detected hit; larger values indicate greater structural similarity to the reference panel. Duplicate structure identifiers were made unique before evaluation.

The final AMPHORA-OS Stage 4 structure panel used the matched full-condition evaluation set, sampled 1,536 generated peptides and 1,259 reference peptides, and retained source records for reproducibility. This panel was compared with the no-condition Stage 3 pair-support structure panel, generated with target-class, genome and text conditions disabled. Because Stage 3 and Stage 4 used different generation contexts and reference panels, this comparison is a lineage-level structure check rather than a matched distribution benchmark. Novel-fold fraction was used as a structure-screening proxy. Because short peptides can be flexible, structure prediction may be uncertain and Foldseek novelty depends on the reference panel and threshold.

#### Statistical analysis and reproducibility

Condition-control point estimates in Fig. 2e and Supplementary Tables 6 and 11 were computed directly from the final Stage 4 partial-condition and shuffled-condition evaluation summaries.

Reported contrasts used unrounded source values for target class plus genome versus target class only, target class plus text versus target class only, matched full versus target class only and matched full versus global shuffled controls. Same-noise directionality intervals were computed from the paired counterfactual rows by resampling paired examples. These summaries quantify evaluation-sample variation for computational readouts and do not replace repeated training seeds.

### Supplementary Tables

| Stage | Main role | Trainable components | Frozen or cached components |
|---|---|---|---|
| Stage 1 | Peptide–genome–text alignment | LoRA peptide adapter; genome/text projection heads | Cached genome and text features |
| Stage 2 | Decodable peptide latent compression | Latent projector; peptide decoder; preservation head | smORFLLM peptide backbone |
| Stage 3 | Unconditional latent-flow backbone selection | Flow backbone; decoder heads for each variant | Target-class, genome and text conditions disabled |
| Stage 4 base | Aligned-context generation | Condition builder; selected flow layers; target-class head | Real-AMP-LoRA-derived initialization; Stage 1 caches; decoder/alignment heads |
| Stage 4 OS | Orthogonal-slot condition evaluation | Selected flow route; separate target-class, genome and text slots; target-class head | Real-AMP-LoRA-derived initialization; rebuilt genome/text condition embeddings |

**Supplementary Table 1. Training scope across AMPHORA stages.** Trainable, frozen and cached components are separated by stage. Full optimization settings are given in the stage-specific Supplementary Methods subsections.

| Encoder configuration | Comparison role | Steps | Avg F1 | Δ Avg F1 |
|---|---|---|---|---|
| ESM2-150M frozen-encoder reference | Protein-LM reference | – | 0.645 | reference |
| smORFLLM selected checkpoint | Peptide-specialized backbone used in AMPHORA | 150K | 0.739 | +0.094 vs ESM2-150M |
| smORFLLM full sparse-fusion recipe | Reduced-budget ablation reference | 60K | 0.717 | reference |
| smORFLLM without sparse biophysical fusion | Sparse-fusion removal ablation | 60K | 0.694 | -0.023 vs 60K full |
| smORFLLM with standard residual pathway | Residual-pathway replacement ablation | 60K | 0.718 | +0.000 vs 60K full |

**Supplementary Table 2. smORFLLM PepBERT9 frozen-transfer and architecture-ablation checks.** All rows were evaluated on nine PepBERT9-style peptide classification tasks with frozen encoder embeddings, mean pooling, linear probes, probe seeds 42, 43 and 44, probe batch size 512 and a short-peptide cap of 50 residues. The selected 150K smORFLLM checkpoint is compared with the ESM2-150M reference under the same protocol. The reduced-budget 60K rows isolate architecture choices within smORFLLM; their deltas are relative to the 60K full sparse-fusion recipe. These checks document peptide-representation accessibility and module sensitivity and are not used as direct AMP-generation endpoints.

| Target class | Train pairs | Val pairs | Train unique | Val unique |
|---|---|---|---|---|
| Bacterium | 73,500 | 3,842 | 13,953 | 775 |
| Fungus | 13,745 | 833 | 5,659 | 328 |
| Virus | 5,999 | 215 | 3,906 | 166 |
| Parasite | 590 | 20 | 361 | 17 |

**Supplementary Table 3. Clean aligned conditional dataset.** The parasite split is substantially smaller than the other target classes. Unique counts refer to distinct peptide sequences within each target-class split.

| Model family | PPL | FPD | pLDDT | $CAMP_{R4}$ | Novel fold | PRDC Prec. | PRDC Rec. |
|---|---|---|---|---|---|---|---|
| Sequence-only flow | 17.339 | 6.872 | 52.812 | 0.323 | 0.023 | 0.876 | **0.538** |
| Global-structure latent flow | 18.092 | 7.346 | 53.014 | 0.350 | 0.022 | 0.882 | 0.479 |
| Joint sequence–structure latent flow | 17.432 | **6.844** | 53.335 | 0.364 | 0.026 | **0.887** | 0.501 |
| Pair-support dual-stream latent flow | **17.166** | 6.987 | **54.633** | **0.438** | **0.032** | 0.871 | 0.508 |

**Supplementary Table 4. AMPHORA model-family comparison.** The table reports the four retained model families: sequence-only flow, global-structure latent flow, joint sequence–structure latent flow and pair-support dual-stream latent flow. The pair-support dual-stream model was selected for conditional generation because it gave the strongest sequence-likelihood and AMP-predictor compromise while retaining competitive structural proxies. PPL is pseudo-perplexity. FPD is peptide feature-distribution distance. $CAMP_{R4}$ is majority-vote AMP predictor support. Novel fold is the Foldseek-derived structural-distance fraction. PRDC precision and recall are local fidelity and coverage readouts from the same Stage 3 evaluation protocol.

| Model | Pred. APEX MIC | APEX pred. <100 $\mu$M | HydrAMP pMIC | Macrel P(AMP) | FPD-ESM2 | MMD | PRDC Prec. | PRDC Rec. | Unique |
|---|---|---|---|---|---|---|---|---|---|
| AMPHORA pair-support fixed-25aa | **302.1** | **0.070** | 0.470 | 0.669 | 8.350 | 0.0070 | **0.934** | 0.403 | **1.0000** |
| AMP-Diffusion-style fixed-25aa | 390.3 | 0.022 | 0.339 | 0.667 | **4.058** | **0.0027** | 0.847 | **0.731** | 0.9982 |
| PepCVAE | 419.5 | 0.008 | 0.201 | 0.619 | 10.157 | 0.0086 | 0.709 | 0.569 | **1.0000** |
| HydrAMP | 348.8 | 0.048 | **0.516** | 0.627 | 11.332 | 0.0088 | 0.601 | 0.468 | **1.0000** |
| AMPGAN | 326.7 | 0.053 | 0.346 | **0.684** | 9.627 | 0.0086 | 0.657 | 0.381 | 0.9996 |
| Training reference | 317.2 | 0.090 | 0.545 | 0.743 | 4.750 | 0.0036 | 0.513 | 0.912 | 1.0000 |

**Supplementary Table 5. Fixed-25aa computational prioritization comparison.** The AMPHORA pair-support and AMP-Diffusion-style fixed-25aa rows are from-scratch training runs on the same 40%-identity-clustered real-AMP split; the AMPHORA row uses the selected pair-support design but does not inherit predicted-AMP Stage 3 weights or use LoRA. Predicted APEX MIC, FPD-ESM2 and MMD are lower-is-better. APEX pred. <100 $\mu$M is the fraction with mean APEX-predicted MIC below 100 $\mu$M. HydrAMP pMIC, Macrel P(AMP), PRDC precision, PRDC recall and uniqueness are higher-is-better. Bold indicates the best generator row for each metric. The training-reference row is an anchor and is not ranked as a generator.

| Condition | Macro F1 | Δ class-only | Bac. rec. | Fung. rec. | Vir. rec. | Par. rec. (n.p.) |
|---|---|---|---|---|---|---|
| AMPHORA-OS full condition | **0.391** | +0.056 | 0.617 | 0.365 | **0.592** | 0.000 |
| Target class only | 0.335 | 0.000 | 0.575 | 0.343 | 0.434 | 0.000 |
| Target class + genome | 0.373 | +0.037 | **0.635** | 0.338 | 0.528 | 0.000 |
| Target class + text | 0.379 | +0.044 | 0.575 | **0.367** | 0.577 | 0.000 |

**Supplementary Table 6. AMPHORA-OS partial-condition comparison.** Full orthogonal-slot conditioning produced the strongest macro F1 in the final Stage 4 evaluation. Macro F1 and recall are shown here; balanced accuracy is displayed in Fig. 2c. All metrics are computed from k-nearest-neighbour target-class assignments in peptide embedding space across the four prespecified target classes. Parasite recall is shown numerically and marked n.p. because validation support was limited. Shuffled-condition controls in Supplementary Table 11 show that aligned genome/text contexts improve over global shuffling in this class-level readout. Fig. 3 separately tests species-aware enrichment of matched generated pools and adds a fresh 7-species APEX post-hoc prioritization readout.

| Species | Manifest rows | Unique seq. | Unique cond. | Reference entries | Prototypes |
|---|---|---|---|---|---|
| *S. aureus* | 19,487 | 11,645 | 24 | 1,000 | 10 |
| *B. subtilis* | 3,051 | 2,625 | 7 | 1,000 | 10 |
| *S. epidermidis* | 2,195 | 1,874 | 4 | 1,000 | 10 |
| *E. faecalis* | 1,941 | 1,406 | 14 | 1,000 | 10 |
| *E. coli* | 15,112 | 8,597 | 26 | 1,000 | 10 |
| *P. aeruginosa* | 9,269 | 5,316 | 16 | 1,000 | 10 |
| *K. pneumoniae* | 2,537 | 1,742 | 11 | 1,000 | 10 |
| *S. enterica* | 3,765 | 2,188 | 18 | 1,000 | 10 |
| *C. albicans* | 8,634 | 5,275 | 7 | 1,000 | 10 |
| *N. glabratus* | 291 | 230 | 2 | 1,000 | 10 |
| *C. tropicalis* | 366 | 254 | 2 | 1,000 | 10 |
| *A. fumigatus* | 201 | 173 | 2 | 1,000 | 10 |

**Supplementary Table 7. Reference sets for the 12-species enrichment benchmark.** Manifest rows and unique sequence counts were computed from the combined train and validation conditional manifests after restricting to the 12 benchmark species. The reference readout sampled 1,000 entries per species after within-species sequence de-duplication; species with fewer than 1,000 unique sequences were sampled with replacement. Multi-prototype assignment used up to 10 MiniBatchKMeans prototypes per species.

| Species | APEX strain outputs | APEX pred. <100 $\mu$M | Mean pred. $\log_{10}$ MIC | Pred. median MIC | Δ vs class-only |
|---|---|---|---|---|---|
| *A. baumannii* | ATCC 19606 | 0.528 | 1.908 | 86.6 | -0.128 |
| *E. faecium* | VRE ATCC 700221 | 0.387 | 2.028 | 158.8 | -0.059 |
| *E. coli* | ATCC 11775; AIC221; AIC222 | 0.386 | 2.092 | 141.0 | -0.103 |
| *P. aeruginosa* | PA01; PA14 | 0.241 | 2.244 | 200.9 | -0.135 |
| *K. pneumoniae* | ATCC 13883 | 0.019 | 2.520 | 375.3 | -0.013 |
| *S. aureus* | ATCC 12600; MRSA BAA-1556 | 0.018 | 2.531 | 389.6 | -0.031 |
| *E. faecalis* | VRE ATCC 700802 | 0.00 | 2.642 | 466.7 | -0.006 |

**Supplementary Table 8. Fresh 7-species APEX post-hoc readout.** The 11 APEX strain-level outputs were grouped into seven species-level target sets as listed. For species with multiple strain outputs, target-specific $\log_{10}$ MIC is the arithmetic mean of the corresponding strain-level predicted $\log_{10}$ MIC values. APEX pred. <100 $\mu$M is the fraction of matched generated peptides with species-level target-specific APEX-predicted MIC below 100 $\mu$M. Mean predicted $\log_{10}$ MIC and predicted median MIC are computed from scored matched generations after this species-level aggregation. Δ vs class-only is matched minus target-class-only mean predicted $\log_{10}$ MIC; negative values indicate lower predicted MIC for matched generation. APEX predictions are computational prioritization readouts and are not wet-lab MIC measurements.

| Metric | Stage 3 no condition | Stage 4 AMPHORA-OS | Stage 4 minus Stage 3 |
|---|---|---|---|
| Mean pLDDT | 54.633 | 67.953 | +13.320 |
| Disorder fraction | 0.414 | 0.099 | -0.315 |
| Contact density | 0.135 | 0.355 | +0.220 |

**Supplementary Table 9. Lineage-level structure check.** The Stage 3 row is the no-condition pair-support panel and the Stage 4 row is the matched full-condition AMPHORA-OS panel. This comparison tests whether the conditional Stage 4 lineage retained predicted structural support relative to its no-condition predecessor. It is separate from the generated-versus-reference structure comparison shown in Fig. 3e.

| Metric | Generated peptides | Reference peptides |
|---|---|---|
| Mean pLDDT | 67.953 | 73.719 |
| Sequence mean pLDDT $\geq$ 70 | 0.430 | 0.629 |
| Contact density | 0.355 | 0.376 |

**Supplementary Table 10. Source metrics for Fig. 3e generated-versus-reference structural support.** The generated group contains 1,536 matched AMPHORA-OS peptides from the final Stage 4 structure panel, and the reference group contains 1,259 curated reference peptides from the corresponding matched full-condition evaluation set. Sequence mean pLDDT $\geq$ 70 is the fraction of peptide structures with mean per-sequence pLDDT at least 70.

| Formal control | Macro F1 | Bal. acc. | KNN acc. | Exact novelty |
|---|---|---|---|---|
| Matched AMPHORA-OS | 0.391 | 0.393 | 0.507 | 0.9997 |
| Global shuffled condition | 0.270 | 0.285 | 0.378 | 0.9990 |
| Within-target-class shuffled condition | 0.381 | 0.384 | 0.495 | 0.9997 |
| Target class only | 0.335 | 0.338 | 0.439 | 0.9997 |
| Target class + genome | 0.373 | 0.375 | 0.485 | 0.9997 |
| Target class + text | 0.379 | 0.380 | 0.489 | 0.9997 |

**Supplementary Table 11. AMPHORA-OS formal condition-control analyses.** All rows use the same ESM2-kNN target-class assignment proxy and are reported as computational class-alignment readouts for condition-control analysis. The table reports the final Stage 4 controls most relevant to the paper claim: partial conditions and genome/text shuffling.
